# PharmCast: rapid generation of three-dimensional pharmacophore fingerprints from two-dimensional structure without conformer generation

**DOI:** 10.64898/2026.09.02.748999

**Authors:** Steven M. Muskal, Malcolm J. McGregor

**Affiliations:** Eidogen-Sertanty, Inc. Oceanside, CA 92056; SRI International, Menlo Park, CA 94025

**Keywords:** pharmacophore, fingerprint, scaffold hopping, surrogate model, virtual screening, applicability domain

## Abstract

A three-dimensional pharmacophore fingerprint records the binding features a molecule can present. It is a description of a hand in search of a glove. Because it is defined by presented features instead of two-dimensional structure, it can identify pharmacophoric similarity between structurally distinct compounds, which is what scaffold hopping and non-obvious me-too design require. The descriptor has remained a niche tool because its cost is dominated by conformer generation. In the reference pipeline, generating 100 conformers requires 2.82 s of the 2.86 s needed to fingerprint one screening collection compound; the bit calculation requires 0.039 s. We therefore removed the conformational stage. PharmCast is a feedforward neural network that predicts all 10,549 bits of a PharmPrint ensemble fingerprint directly from a SMILES string. On the same machine, PharmCast generated pharmacophore fingerprints for two molecules and compared them in 0.584 ms, whereas the conventional conformer-based pipeline took 5.71 s. PharmCast version 10 was trained on 5,887,229 molecules drawn from a screening collection, activity-backed ChEMBL compounds from 142 to 1000 Da, and peptide loops excised from crystal structures. We evaluated 155,648 purchasable catalog compounds excluded from every training set, 139,700 activity-backed ChEMBL compounds not present in the version 10 training set, and 13,500 peptide loops reserved for testing. Median fingerprint error, Pearson r, and pairwise ranking accuracy were 0.008, 0.980, and 0.936 for screening collection chemistry; 0.016, 0.984, and 0.952 for loop peptides; and 0.027, 0.936, and 0.889 for activity-backed ChEMBL compounds. The reference calculation reproduces itself at an error of 0.006 and r of 0.995. Ensemble pharmacophore fingerprints can therefore be predicted from two-dimensional structure alone at a cost suitable for large-scale collection screening and virtual library exploration.

## **1.** Introduction

### 1.1 A century of the pharmacophore

The notion that binding depends on a small set of chemical features held in a particular spatial arrangement, and not on the skeleton carrying them, is among the oldest working ideas in medicinal chemistry. It is generally credited to Paul Ehrlich, whose 1909 account of chemotherapy illustrates the principle that a compound must possess both a group conferring affinity and a group conferring action.^1^ Güner and Bowen have traced the concept to Ehrlich’s late nineteenth century work, its movement toward a modern spatial definition through Schueler in 1960,^2^ and its development into pharmacophore modeling by Kier from 1967,^3^ who described the common features of ligands for central nervous system receptors and named the muscarinic pharmacophore.^4^

What made the idea computable was the ability to search three dimensional structures instead of reasoning about them by hand. Gund framed the problem of three dimensional pharmacophoric pattern searching,^5^ and by the late 1980s systems such as 3DSEARCH^6^ could take a query of typed features and interfeature distances and retrieve every molecule in a database able to present it. A two-dimensional substructure query asks what a molecule is made of; a pharmacophore query asks what it can present; and the two questions have different answers. Asking the second question at scale has never been practical.

### 1.2 From query to descriptor

A query returns a list, whereas a descriptor returns a number for every molecule, and it is the descriptor that diversity analysis, similarity searching and quantitative modeling all require. The step from one to the other was to stop asking whether a molecule matches a single pharmacophore and to begin recording which of all possible pharmacophores it can present.

Pickett, Mason and McLay took that step with pharmacophore derived queries,^7^ profiling libraries by the three-point pharmacophores their members express. The present authors then defined the fingerprint used here^8,9^: a fixed-length bit string in which each bit represents one three-point pharmacophore: a triple of feature types and the three binned distances between them. A molecule sets a bit when any accessible conformation presents that triangle. Mason and colleagues subsequently extended the construction to four-point pharmacophores,^10^ which resolve the chirality that three-points cannot.

The defining property of the descriptor is that it is nearly blind to scaffold. Figure 2 makes the point with the example from the original paper. Estradiol and diethylstilbestrol are both estrogen receptor ligands, they share a two-dimensional Morgan similarity of only 0.16, and they nevertheless present an acceptor, a donor and an aromatic ring at three distances falling in the same three bins; they therefore set the same bit. This is what scaffold hopping looks like from inside the descriptor, and it is why topological pharmacophore searching became a recognized route to it.^11,12,13^

**Figure 1.**
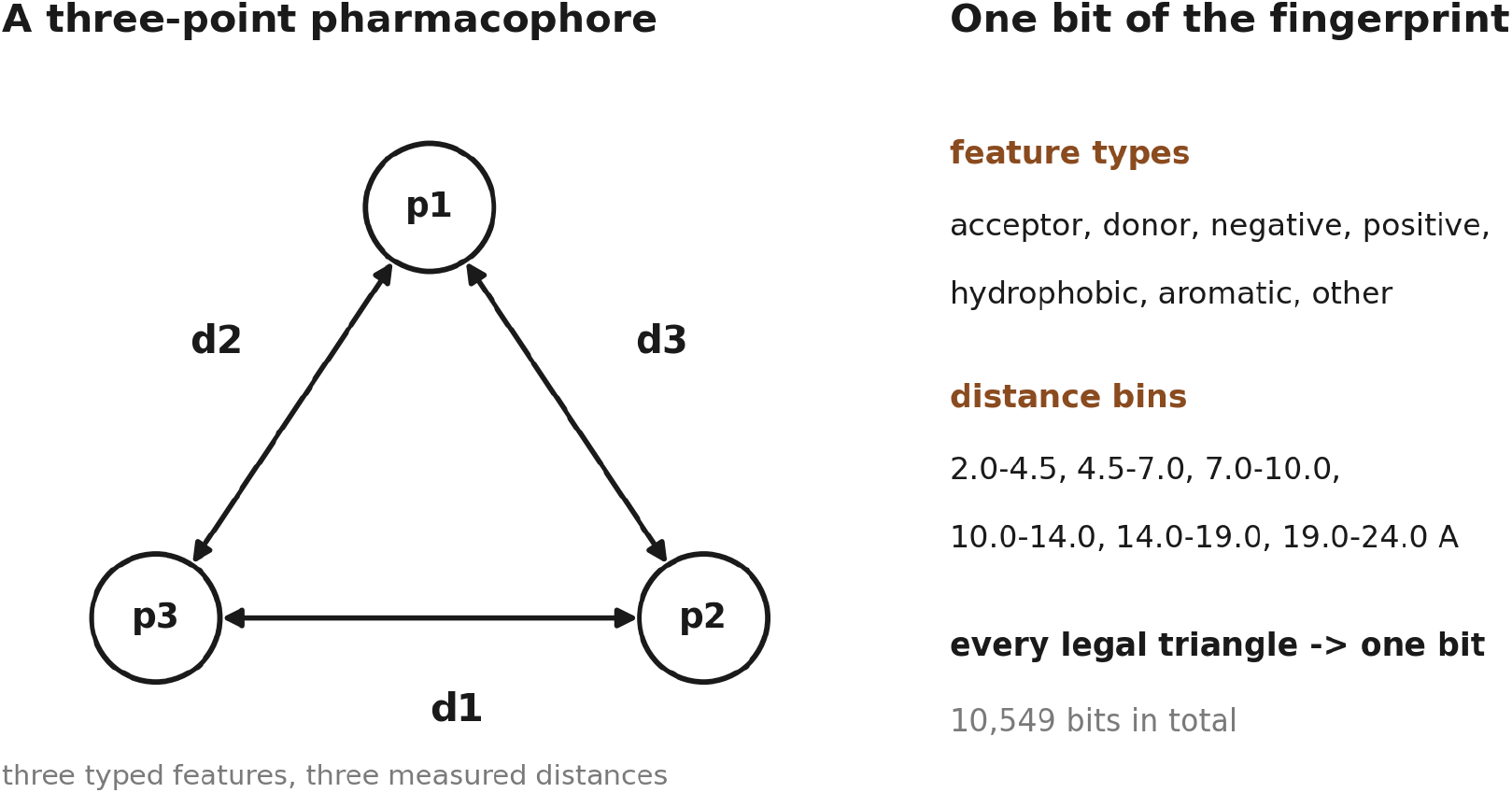
What one bit is. A three-point pharmacophore has three typed features and the three binned distances between them; enumerating every triangle that satisfies the triangle inequality on the bin bounds gives 10,549 distinct pharmacophores. Reproduced from McGregor and Muskal.^8^

**Figure 2.**
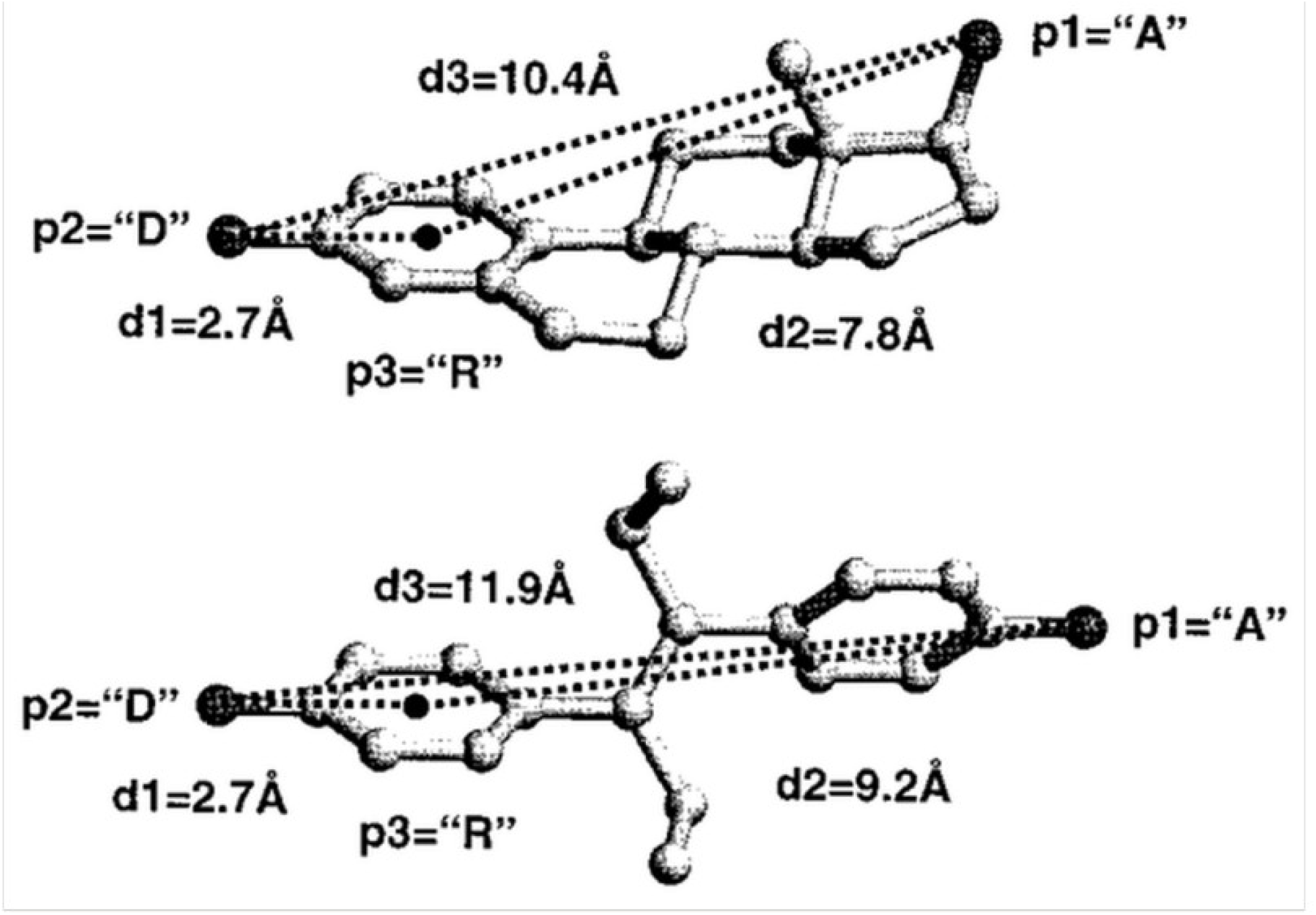
The same pharmacophore on two unrelated scaffolds. Estradiol (above) and diethylstilbestrol (below) each present an acceptor (p1), a donor (p2) and an aromatic ring (p3). The distances differ, 2.7, 7.8 and 10.4 Å against 2.7, 9.2 and 11.9, but all three fall in the same bins, so both molecules set the same bit. Reproduced from McGregor and Muskal.^8^ Regenerating the geometry independently from a 100 conformer ETKDGv3 ensemble gives 2.8, 8.2 and 10.9 Å for estradiol and 2.8, 9.3 and 12.1 for diethylstilbestrol, the same three bins. These molecules have a low 2D similarity of 0.16 by Morgan fingerprints, but a relatively strong pharmacophoric similarity of 0.72 by the original conformation-dependent 3D method, and 0.78 by PharmCast version 10.

Pharmacophore methods have since become standard in structure based form deriving models from protein bound ligands^14^ and in ligand based form throughout virtual screening and library design. Leach, Gillet, Lewis and Taylor give a thorough account of the field as it stood.^15^

### 1.3 Computational cost

The method carries one structural cost. A molecule does not have a single three-dimensional shape, so the fingerprint is defined over a conformer ensemble, conventionally the bitwise OR across many conformers. Generating that ensemble dominates the computational cost.

Timed on screening collection compounds with the pipeline used here, the full route requires 2.86 s per molecule, of which 2.82 s is conformer generation with RDKit ETKDGv3 followed by UFF minimization and 0.039 s is the fingerprint calculation. At that rate a five million compound collection is roughly five and a half core months. Conformer generation is itself a mature and carefully engineered problem, whether by knowledge augmented distance geometry^16^ or by systematic torsion driving,^17^ and its cost is not waste; it is what the answer requires.

The consequence has been that pharmacophore similarity is confined to small, curated sets. It is not usable as a screening objective over the full screening collection, nor inside an optimization loop, which is precisely where a three dimensional similarity measure would be most valuable. Two-dimensional fingerprints such as extended-connectivity fingerprints^18^ are used instead because they are computationally affordable, although they answer a different question. Since the fingerprint calculation is only 1.7% of the total, accelerating it changes nothing; anything that materially changes the economics must remove the conformational stage, not speed it up.

### 1.4 This work

We predict the ensemble fingerprint directly from two-dimensional structure and omit conformer generation. An ensemble fingerprint is an OR over conformations, so a bit records that a molecule can present a given triangle in some accessible conformation, without asserting that it presents it in any particular pose. That is closer to a property of molecular constitution than of any single geometry, and constitution is what a two-dimensional structure encodes.

PharmCast is deliberately simple: a feedforward network maps a circular fingerprint to all 10,549 pharmacophore bits. More elaborate learned representations, including graph convolutional^19^ and message-passing^20^ networks, may improve performance.

Figure 2 illustrates the distinction between pharmacophoric and two-dimensional similarity. Structurally distinct ligands for the same target are common enough to motivate scaffold hopping^11,12,13^. Böhm, Flohr, and Stahl note that changing a central template can yield a patentable structure and that pharmacological and legal definitions of similarity differ^12^. A screening-collection-scale pharmacophore fingerprint can therefore support scaffold hopping and prioritization of structurally novel compounds. This work provides (i) a surrogate model and a measurement of its agreement with the reference calculation relative to that calculation’s reproducibility; (ii) an explicit applicability domain that quantifies failure and relates it to training-set composition; and (iii) a version series showing that additional data from the same chemical space did not repair out-of-domain error, whereas adding the missing chemistry did.

## 2. Methods

### 2.1 The fingerprint

Figure 1 defines one bit. Each fingerprint contains 10,549 bits, one per three-point pharmacophore. A bit contains three feature types drawn from acceptor, donor, negative, positive, hydrophobic, aromatic and other, together with the three pairwise distance bins between them. Distance bins are 2.0 to 4.5, 4.5 to 7.0, 7.0 to 10.0, 10.0 to 14.0, 14.0 to 19.0 and 19.0 to 24.0 Å. The enumeration is deterministic: distances are sorted ascending with ties broken by feature type so that each triangle has one canonical form, combinations violating the triangle inequality on the bin bounds are discarded, and the survivors are numbered sequentially. This encoding is the three-point PharmPrint, and a similarity comparison between two of them is a PharmSim score.

Reference fingerprints were generated identically for each source set, i.e. 100 conformers from isomeric SMILES using RDKit ETKDGv3 with a fixed seed followed by UFF minimization, largest fragment, hydrogens added, and per conformer fingerprints combined by bitwise OR into one ensemble record. Using a single recipe throughout is what permits scores to be compared across source sets.

### 2.2 Training data

PharmCast version 10 uses three data sources: the Enamine screening collection, activity-backed ChEMBL compounds, and protein loop peptides. The June 2026 Enamine screening collection held 4,774,670 source compounds, of which 4,612,044 survived property filtering. Median molecular weight is 344 Da (range 116 to 599; 5th and 95th percentiles 255 and 465). Compounds have median values of 24 heavy atoms, 3 rings (2 aromatic), cLogP 2.7, and polar surface area 71 Å²; 8% have defined stereochemistry. Compounds in the file are labeled by Enamine as HTS, Advanced, Legacy or Functional, of which only HTS is the actively stocked screening library, a distinction that matters whenever a screening result is quoted as purchasable.

We selected the three sources because (i) the screening collection contains synthesized molecules intended to interrogate biology; (ii) activity-backed ChEMBL contains compounds that have been synthesized and tested against biological endpoints, demonstrating synthetic accessibility and potential biological relevance; and (iii) protein loop peptides come from regions of X-ray structures outside regular secondary structure, where loops often participate in antigen recognition, protein-protein interactions, or protein-ligand interactions.

The collection is diverse by scaffold and redundant by similarity, and both properties matter. In a 200,000 molecule sample there are 135,768 distinct scaffolds, 88% of which appear exactly once, and the ten commonest scaffolds together hold 4.0% of the collection. The median compound nevertheless has a nearest neighbor at 0.714 Tanimoto on Morgan fingerprints, 54.9% have a neighbor at 0.70 or closer, and 64,473 compounds have a two-dimensional identical twin, so a raw compound count overstates how much distinct chemistry is present.

The full peptide set contains 136,494 loops of two to six residues excised from Protein Data Bank crystal structures^21^, all from structures at 2.5 Å resolution or better and R-free 0.23 or better, and capped with the flanking backbone atoms present in the structure: the preceding residue’s C and O as an acetyl and the following residue’s N and CA as an N-methylamide. Each peptide fingerprint is the bitwise OR of 100 ETKDGv3 conformers, each optimized with UFF and converted to a reference fingerprint, together with every rigid crystal conformation observed for that peptide.

Protein loop peptides extend the conformational range represented in the training set. A surrogate for an ensemble fingerprint is hardest where a molecule has the most conformational freedom, and screening collection chemistry holds very little of it; the median loop has 16 rotatable bonds, compared with 5 for a screening-collection compound. We filled that range with loops excised from deposited structures, each a fragment observed in a protein structure. Median bit count is 412 pharmacophores per fragment against 534 for a collection compound.

For the peptide holdout, 13,500 loops, 9.9% of the full peptide set, were reserved for testing and never used for training. The holdout was stratified by sequence length from two to six residues, molecular-weight quintile, and the number of observed crystal conformations in four bands: 1, 2 to 3, 4 to 10, and more than 10. Every stratum contributed 9.9%, and the holdout median molecular weight matches the training set at 576.6 Da. The 13,500 held-out loops and the 122,994 training loops together account for all 136,494. Their reference fingerprints were generated by one procedure, so the only difference between them is whether a loop was available during training.

The ChEMBL set includes every ChEMBL^22^ molecule from 142 to 1000 Da with at least one recorded activity, establishing that the compound was synthesized, purified, and assayed. Fingerprinting proceeds in ascending molecular weight so that coverage extends incrementally from chemistry already represented in the training set. Of these molecules, 1.6% fail to embed.

Each source set is sampled to the same size, 6,000 molecules, because nearest neighbor similarity rises with sample size and unequal samples cannot be compared. These values are therefore comparable to one another but not to a figure computed over a whole source set: the full 4.6 million compound collection has a median nearest neighbor of 0.714 against 0.389 in a 6,000 molecule sample of it. That the peptides are the least diverse of the three is expected, since the same loop sequence recurs across many structures.

### 2.3 Model architecture and training

The network takes a 2,048 bit binary Morgan fingerprint of radius 2 alongside 11 whole molecule descriptors, 2,059 inputs in all, and predicts all 10,549 pharmacophore bits through two hidden layers of 1,024 and 512 units, for 8.05 million parameters. The descriptors are molecular weight, cLogP, topological polar surface area, rotatable bonds, rings, aromatic rings, hydrogen bond donors, hydrogen bond acceptors, heavy atoms, fraction of sp3 carbon and Labute accessible surface area. Training minimizes binary cross entropy.

Training and all timing measurements were performed on a single Apple Mac Studio with an M3 Ultra chip, 28 CPU cores, a 60 core GPU and 256 GB of unified memory, running PyTorch 2.9.1. From PharmCast version 8 onward, training used the Metal Performance Shaders backend on the GPU; the timing measurements reported here were made on the CPU, and inference uses no GPU. Each version retained the best set of weights obtained during training.

Saquinavir and indinavir illustrate in Figure 5. Their Morgan similarity is 0.303, consistent with different chemical series, whereas their pharmacophore similarity is 0.841. Both compounds lie in the sparsely represented 600 to 700 Da range. Neither compound occurs in any training set. The two structures are CC(C)(C)NC(=O)[C@@H]1C[C@@H]2CCCC[C@@H]2CN1C[C@@H](O)[C@H](C c1ccccc1)NC(=O)[C@H](CC(N)=O)NC(=O)c1ccc2ccccc2n1 for saquinavir and CC(C)(C)NC(=O)[C@@H]1CN(Cc2cccnc2)CCN1C[C@@H](O)C[C@@H](Cc1ccccc1)C(=O)N[C@H]1c2ccccc2C[C@H]1O for indinavir.

**Figure 3.**
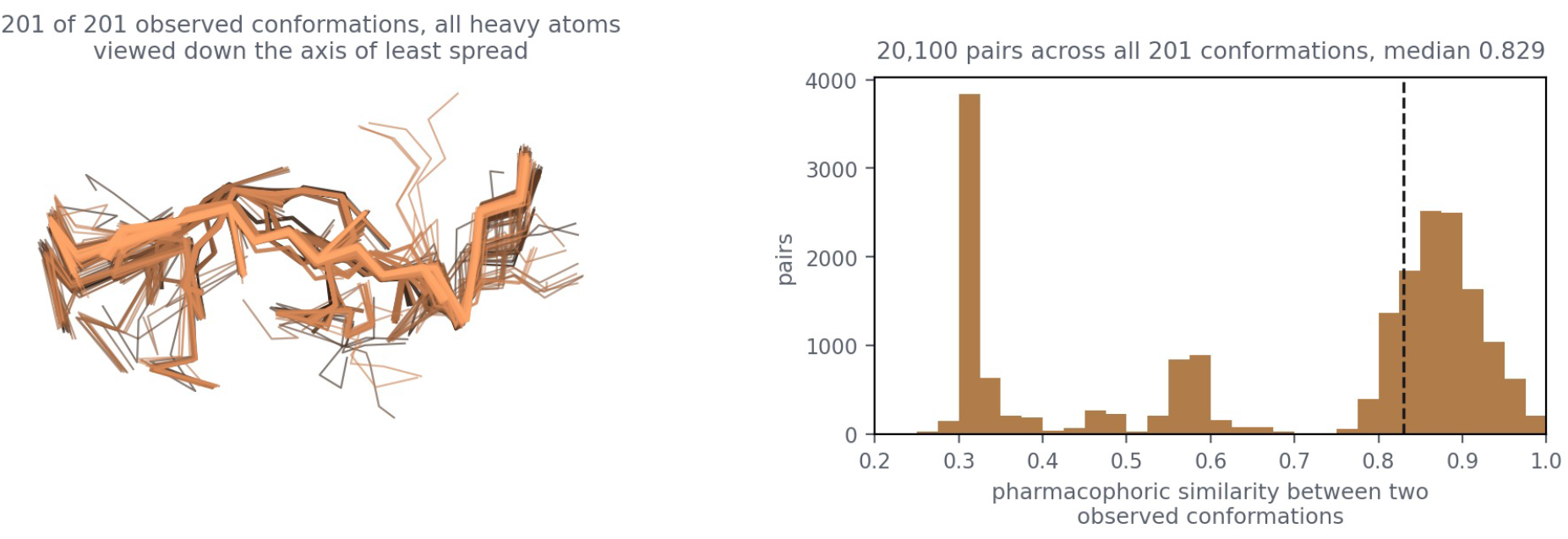
One loop sequence and every deposited conformation of it. The sequence LGGK appears 217 times across 61 Protein Data Bank entries; the 201 instances sharing the same 25 heavy atoms are drawn superimposed on their backbones, viewed down the axis of least spread, with a maximum backbone RMSD of 2.59 Å. Their 20,100 pairwise comparisons have a median of 0.829. Those 20,100 comparisons fall into three families, near 0.31, near 0.58 and a dominant one near 0.87.

**Figure 4.**
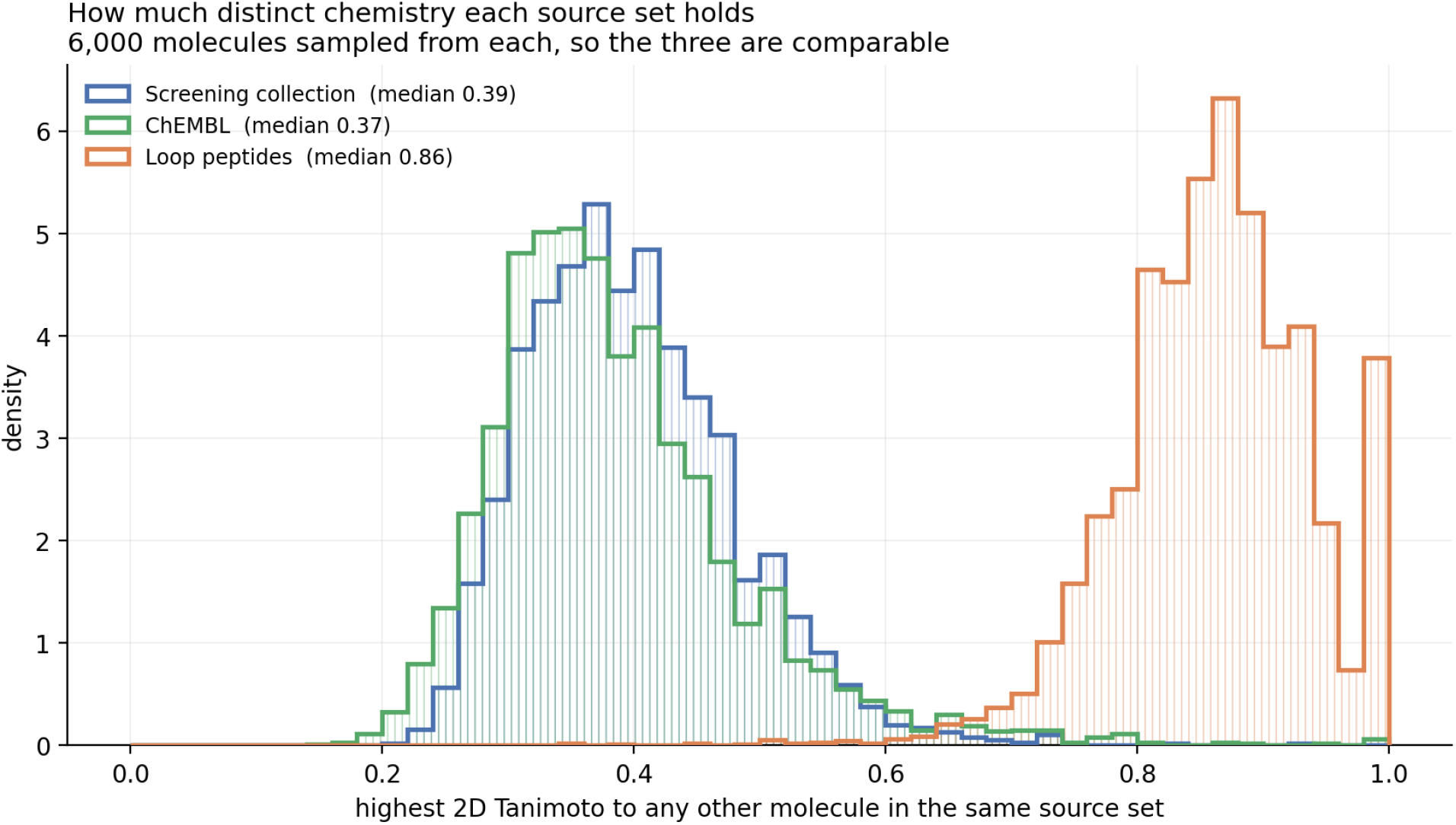
How much distinct chemistry each source set holds in the version 10 training set. Each distribution uses 6,000 molecules, and each value is a molecule’s highest two-dimensional Tanimoto to another molecule in the same source set. Medians are 0.389 for the screening collection, 0.368 for ChEMBL, and 0.864 for loop peptides.

**Figure 5.**
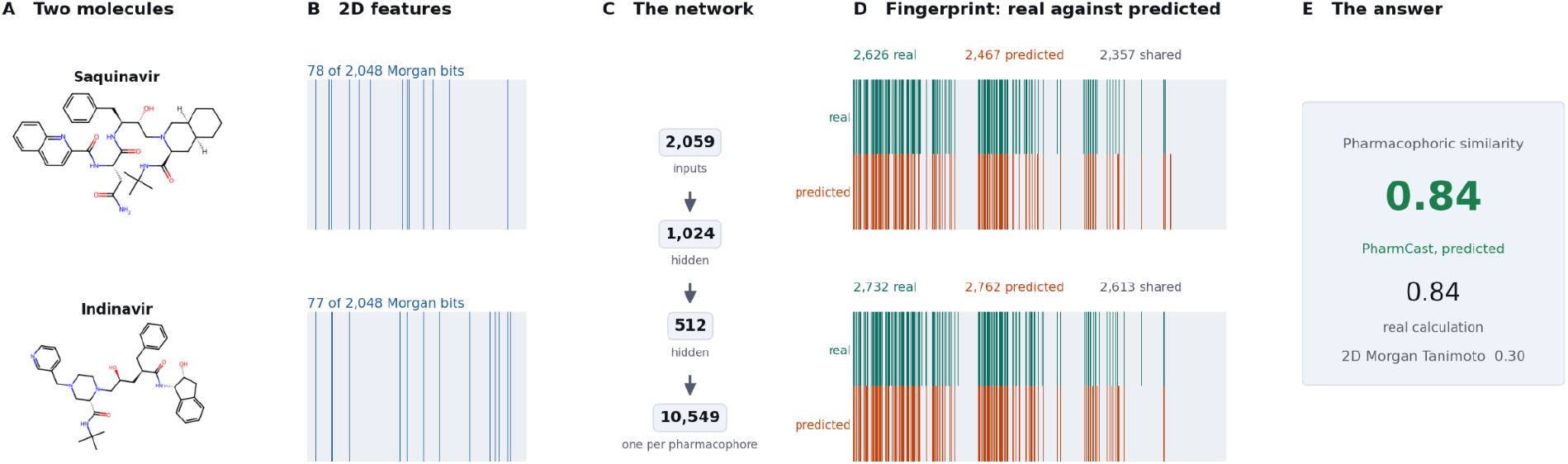
The two routes to a fingerprint, shown for saquinavir and indinavir, HIV-1 protease inhibitors of unrelated scaffold. The conventional route generates a conformer ensemble and applies the reference calculation; PharmCast predicts the ensemble record from two-dimensional structure. PharmCast predicts a pharmacophore Tanimoto of 0.843, against 0.841 from the reference calculation and a two-dimensional Morgan Tanimoto of 0.303.

PharmCast version 10 was trained on 5,887,229 molecules drawn from screening collection compounds, activity-backed ChEMBL compounds from 142 to 1000 Da, and protein loop peptides, with a molecular-weight-stratified stopping set reserved for early stopping; feature normalization was fitted only on the training rows.

The stopping set is distinct from the test sets. Version 10 has three unseen test populations: 13,500 peptides reserved from the full peptide set; activity-backed ChEMBL molecules not present in the version 10 training set; and purchasable Enamine screening-collection compounds excluded by the training-set ingest filter. None was used for training or early stopping.

### 2.4 Test sets and evaluation

Version 10 is evaluated on three chemically distinct populations because pooling would conceal their different failure modes. All are unseen during training.

● Loop peptides: 13,500 molecules, 9.9% of the full peptide set, reserved for testing and never trained on. Sampling was stratified by sequence length from two to six residues, by molecular-weight quintile, and by the number of crystal conformations deposited for that peptide, grouped as one, two or three, four to ten, and more than ten. Each stratum contributed 9.9%; the holdout and the training set have the same median molecular weight, 576.6 Da.
● ChEMBL: 139,700 activity-backed molecules not present in the version 10 training set and held out.
● Screening collection: 157,335 real, purchasable Enamine screening-collection compounds from **the June 2026 Enamine** catalog that the training-set ingest filter excluded. Canonical SMILES were compared with all 4,617,292 screening collection structures, and two matches were removed. Molecular weight spans 42 to 1000 Da. Of these, 155,648 were fingerprinted and scored, and 1,687 could not be embedded.

Evaluation uses all 13,500 peptide holdouts, all 139,700 held-out ChEMBL molecules, and the 155,648 screening collection test cases with reference fingerprints. Pair statistics are estimated from five million pairs drawn from each population, and ranking accuracy from one million molecule triples.

Fingerprint error is the absolute difference between the predicted and the reference Tanimoto for a pair of molecules. Ranking accuracy is measured on triples of one query and two candidates. It is the fraction of triples in which the surrogate ranks the two candidates in the same order as the reference calculation, with exact ties set aside.

Agreement describes a single molecule rather than a pair. It is the Matthews correlation coefficient^23^ between the predicted and the reference fingerprint bits, and it is labeled MCC in every table and figure. A plain accuracy would not serve here, because the fingerprints are sparse and a score over 10,549 bits would be carried by the bits that are zero in both. The pair statistics follow from this per molecule distribution.

A ceiling is required in order to interpret either quantity. Repeating the reference calculation on the same molecules with a different embedding seed gives a fingerprint error of 0.006 at r 0.995, and no surrogate can be expected to do better than the target’s own reproducibility.

Historical version analyses score every released model on the same 5,000 screening collection test cases. Table 10 also reports ranking accuracy on 573 real ChEMBL compounds above 600 Da that were excluded from every model’s training set. The fixed populations allow the models to be compared on one basis.

### 2.5 Reference fingerprint coverage

Table 1 gives the reference fingerprints available when version 10 was trained against the eligible molecules in each source. The ChEMBL remainder is chemistry that later versions can draw on, both as training data and as held-out molecules for testing.

**Table 1.** Reference fingerprint coverage by source when PharmCast version 10 was trained.

| Source | Reference fingerprints | Eligible molecules | Coverage |
| --- | --- | --- | --- |
| Screening collection | 4,609,488 | 4,617,292 | 99.8% |
| ChEMBL | 1,214,214 | 2,737,190 | 44.4% |
| Loop peptides | 136,494 | 136,494 | 100% |
| Total | 5,960,196 | 7,490,976 | 79.6% |

The screening collection contributes 4,609,488 reference fingerprints from 4,617,292 eligible structures. The peptide source contributes 136,494 reference fingerprints, of which 122,994 were used for training and 13,500 reserved for testing. ChEMBL contributes 1,214,214 reference fingerprints from an eligible set of 2,737,190 molecules.

PharmCast version 10 was trained on 5,887,229 molecules from the screening collection, ChEMBL and protein loop peptides, with a molecular-weight-stratified stopping set reserved for early stopping. The test populations are defined in section 2.4.

## 3. Results

All results in this section were obtained with PharmCast version 10. Figures 9 and 10 and Table 10 place it in the version series, where agreement has flattened across versions 8, 9 and 10, which suggests what further growth in the training set is likely to buy..

### 3.1 Computational performance

All timings were measured on one machine, an Apple M3 Ultra with 28 CPU cores, 20 performance and 8 efficiency, a 60 core GPU and 256 GB of unified memory, running macOS 26.5.1 with PyTorch 2.9.1 on CPU. The GPU was not used, and both routes were timed on the same hardware. These are inference timings, and a GPU would not materially change them: the forward pass is 0.009 ms of the 0.288 ms per molecule, so accelerating it cannot move the total. Training is another matter, and there the same GPU is 12.5 times faster per step.

Table 2 reports the end-to-end cost of predicting every fingerprint from SMILES and screening the complete collection: 29.3 min at 2,644 compounds per second on six worker processes, including screening collection input and featurization. Screening a collection whose fingerprints already exist is a bitwise comparison, which costs 0.012 ms per pair. The 3,697 core-hour conventional value was extrapolated from the per-molecule rate because the collection has been fingerprinted by that route only in multiday production runs.

**Table 2.**
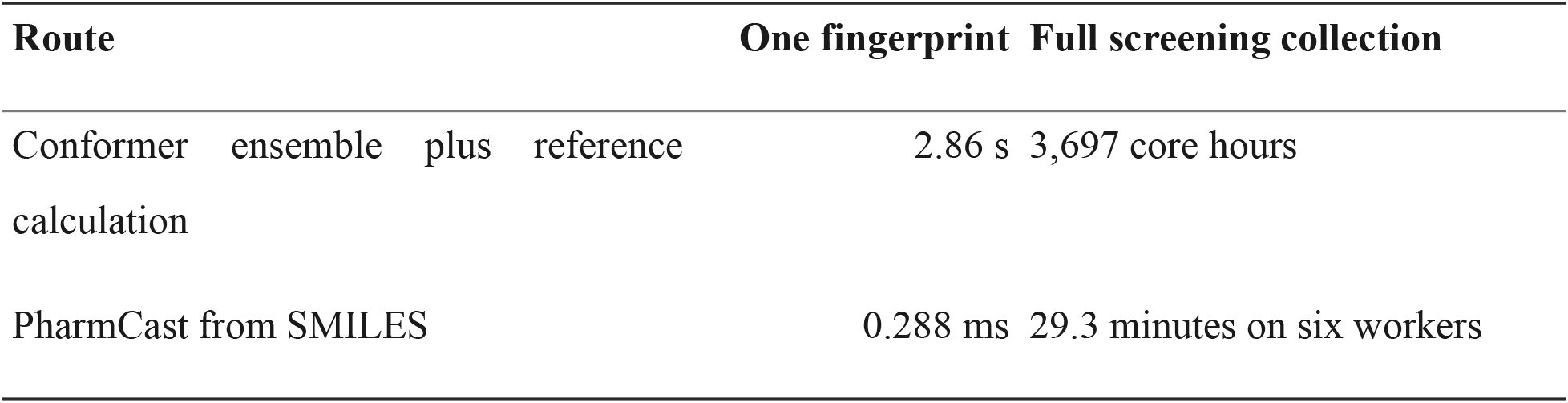
Cost of generating one pharmacophore fingerprint, and of fingerprinting the complete collection of 4,653,831 compounds, by route.

The per molecule figures split as follows. 2.82 s is conformer generation with RDKit ETKDGv3 followed by UFF minimization, and 0.039 s is the fingerprint calculation over the ensemble. PharmCast replaces both with featurization and one forward pass. In a library-scale batch, this takes 0.288 ms per molecule, including 0.009 ms for the forward pass. Processed alone, a molecule takes 0.357 ms, including 0.294 ms for featurization.

Comparing two fingerprints that already exist is a Tanimoto over 10,549 bits and costs 0.012 ms by either route, so the comparison is not where the difference lies. Starting instead from two SMILES strings, PharmCast returns a similarity in 0.584 ms at library scale against 5.71 s for the conventional route. Everything separating those two figures is conformer generation.

### 3.2 Fidelity by test population

PharmCast version 10 was trained on 5,887,229 molecules. It is evaluated on 155,648 Enamine screening collection test cases, 139,700 activity-backed ChEMBL compounds not present in the version 10 training set, and the 13,500-peptide reserved holdout. Median per molecule MCC is 0.881, 0.860, and 0.914 respectively.

### 3.3 Independence from two-dimensional similarity

Figure 8 plots PharmCast’s pairwise similarity against the reference calculation’s, as Figure 6 does, with each pair colored by the two-dimensional Morgan similarity of the two molecules. The points follow the diagonal at every color, so PharmCast tracks the reference calculation whether or not the two molecules are two-dimensionally alike.

**Figure 6.**
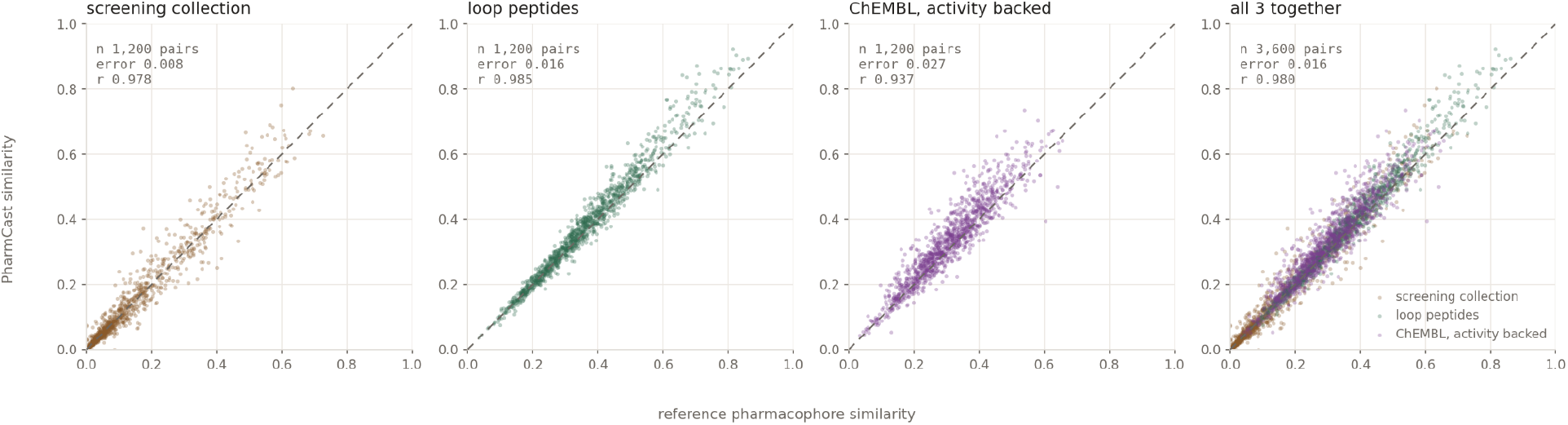
Predicted against real pairwise similarity for PharmCast version 10. Peptides are the reserved holdout, screening collection compounds are Enamine test cases, and ChEMBL compounds were not present in the version 10 training set. Each population panel draws 1,200 pairs at random from 900 uniformly sampled molecules; the combined panel pools 3,600 pairs. Panel error and r use those pairs. The pair count keeps the scatter legible; Table 3 reports all tested molecules. The dashed line is exact agreement. The weakest population is ChEMBL, for which median error is 0.027 and roughly one pair in four falls outside 0.05.

**Figure 7.**
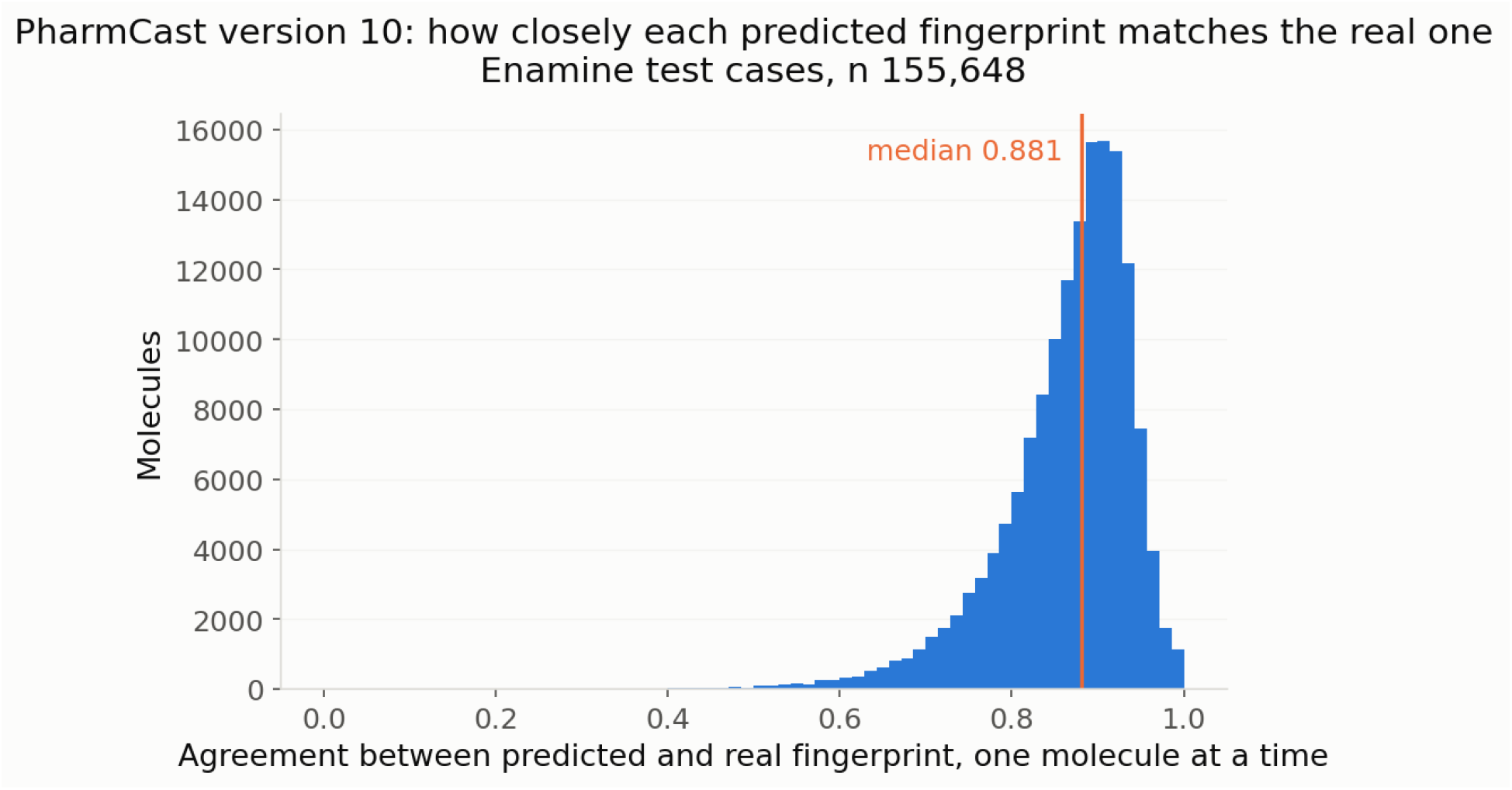
How closely each predicted fingerprint matches the reference, as MCC per molecule, for PharmCast version 10 on 155,648 Enamine test cases. Median 0.881. The pairwise similarity results are a consequence of this distribution.

**Figure 8.**
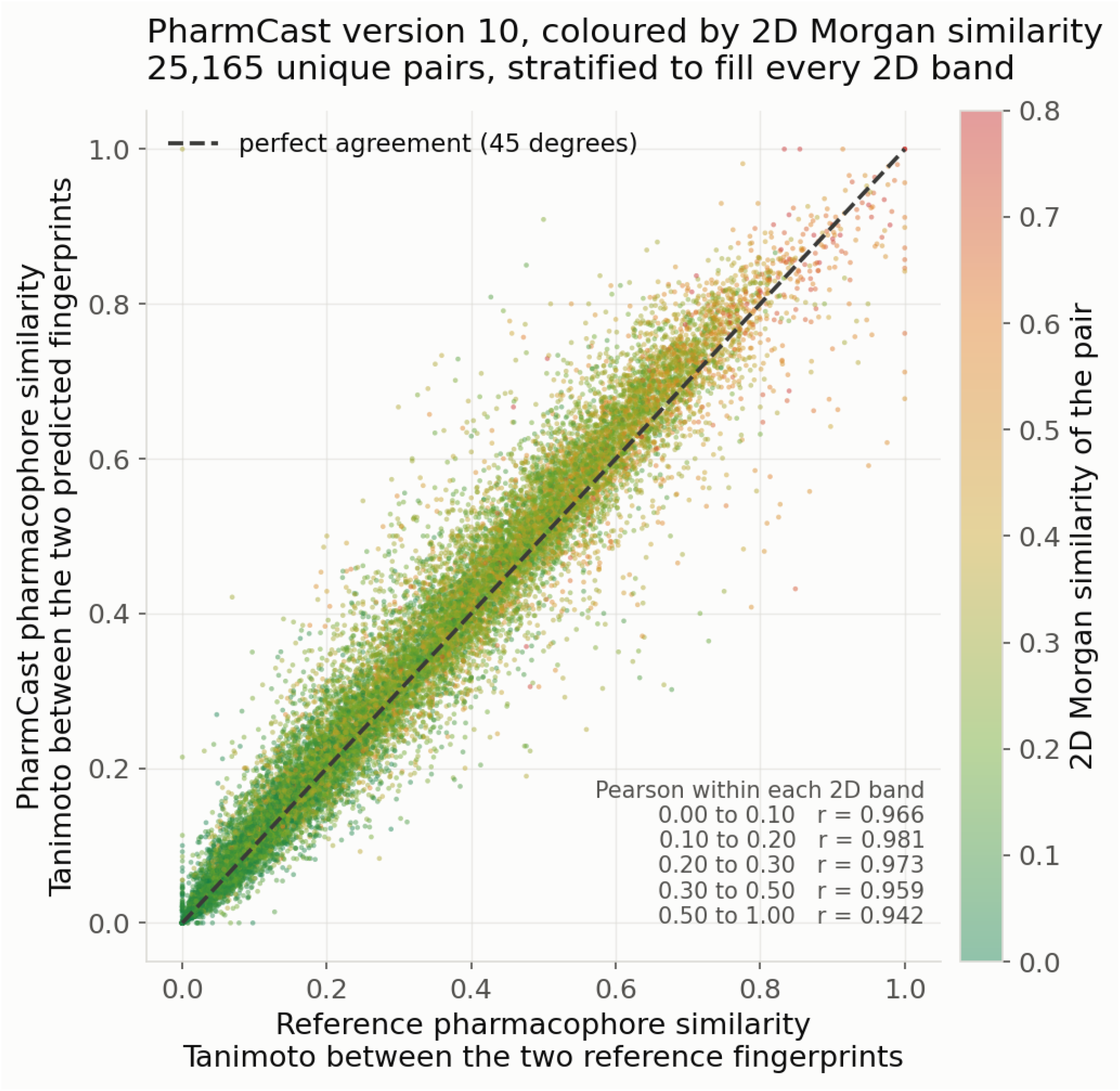
Predicted against real pairwise similarity, colored by two-dimensional Morgan similarity, green for the most dissimilar pairs through red for the least. PharmCast version 10 on 5,000 held-out Enamine test cases. Agreement is flat across the color range, consistent with prediction of three-dimensional feature geometry independently of two-dimensional similarity.

**Table 3.** Fidelity by population for PharmCast version 10, against the reference calculation’s own reproducibility. Screening collection is the Enamine test cases, ChEMBL is activity-backed chemistry not present in the version 10 training set, and loop peptides are the reserved holdout. Median MCC uses every molecule; the pair statistics come from five million pairs sampled per population and the ranking accuracy from one million triples, because the possible pairs grow with the square of the population. The final row compares the reference calculation with itself and gives the reproducibility ceiling.

| Population | Molecules tested | Median MCC | Median error | Pairs within 0.05 | Pearson r | Ranking accuracy |
| --- | --- | --- | --- | --- | --- | --- |
| Screening collection | 155,648 | 0.881 | 0.008 | 89.3% | 0.980 | 0.936 |
| Loop peptides | 13,500 | 0.914 | 0.016 | 88.3% | 0.984 | 0.952 |
| ChEMBL, activity backed | 139,700 | 0.860 | 0.027 | 74.8% | 0.936 | 0.889 |
| Reference self-comparison | N/A | N/A | 0.006 | N/A | 0.995 | Ceiling |

**Table 4.** Composition of the PharmCast version 10 training set by mass, counted before one percent was set aside for early stopping. The three sources cover 74 to 1000 Da between them. The screening collection is the bulk of the training set and is filtered at 600 Da on ingest, so the range above 600 Da is carried by ChEMBL and by the loop peptides.

| Source | Molecules | Share | Mass range | Median |
| --- | --- | --- | --- | --- |
| Screening collection | 4,609,488 | 77.51% | 74 to 600 Da | 344 |
| ChEMBL, activity backed | 1,214,214 | 20.42% | 142 to 1000 Da | 323 |
| Protein loop peptides | 122,994 | 2.07% | 116 to 988 Da | 577 |
| Total | 5,946,696 | 100% | 74 to 1000 Da | 341 |

**Table 5.**
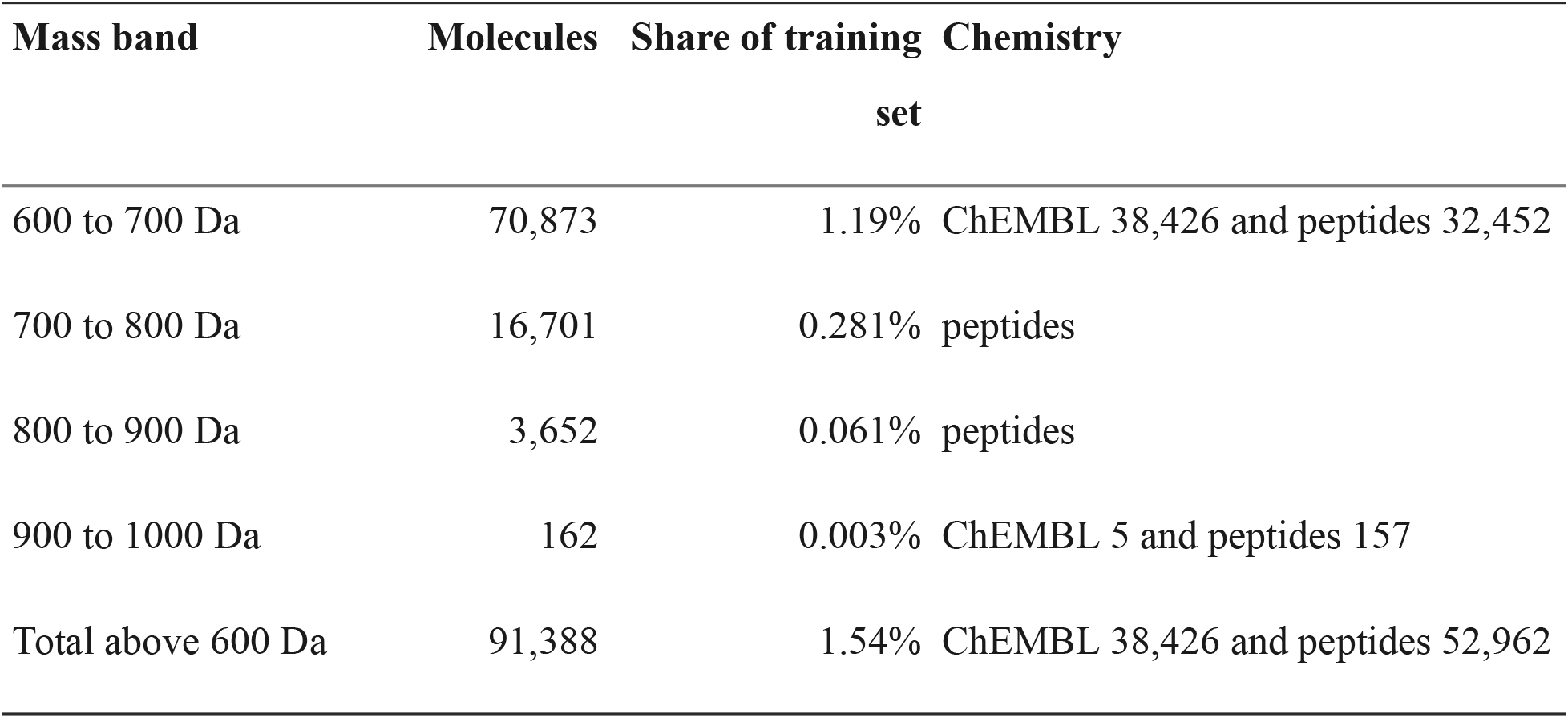
Molecules above 600 Da, by mass band, in the PharmCast version 10 training set. Counts come directly from the training set indexes.

**Table 6.** Per molecule agreement against molecular weight for PharmCast version 10. Screening collection is the Enamine test cases, ChEMBL is the held-out activity-backed set, and loop peptides are the reserved holdout. Cells holding fewer than 20 molecules read not measured.

| MW band | Screening collection | ChEMBL | Loop peptides |
| --- | --- | --- | --- |
| up to 300 Da | 0.872 | not measured | 0.902 |
| 300 to 400 Da | 0.887 | 0.860 | 0.938 |
| 400 to 500 Da | 0.887 | not measured | 0.937 |
| 500 to 600 Da | 0.899 | not measured | 0.922 |
| 600 to 700 Da | 0.874 | not measured | 0.899 |
| 700 to 1000 Da | 0.844 | not measured | 0.867 |

The 5,000 molecules form 12,497,500 pairs. Each was scored for two-dimensional similarity, and the figure draws 6,000 from each of the bands 0 to 0.1, 0.1 to 0.2, 0.2 to 0.3 and 0.3 to 0.5, together with all 1,165 pairs above 0.5, for 25,165 in total. Balancing the bands is necessary because the library is not balanced: in a uniform draw, 97.5% of pairs fall below 0.20, and the two-dimensionally similar end of the panel would be nearly empty. Pearson agreement is 0.966, 0.981, 0.973, 0.959 and 0.942 across those bands, from the least two-dimensionally similar to the most. Scaffold hopping depends on this. A method that recovered only two-dimensional similarity could not find matches across unrelated scaffolds, which is what a pharmacophore fingerprint is for.

### 3.4 Applicability domain

PharmCast version 10 is trained across 74 to 1000 Da. Performance within that span depends on training-set composition.

Screening collection and peptide agreement both vary by mass. Peptide agreement runs from 0.938 at 300 to 400 Da down to 0.867 above 700 Da, and screening collection agreement from 0.899 at 500 to 600 Da down to 0.844 above 700 Da. The held-out ChEMBL molecules span 374.4 to 398.1 Da, so they populate the 300 to 400 Da band alone and measure a narrow mass slice rather than ChEMBL generally; there they read 0.860 against 0.887 for screening collection compounds of the same size. Molecular weight alone therefore does not define the boundary. Training-set coverage and chemical class both matter, and ChEMBL is the most heterogeneous source and the least complete, at 44.4% fingerprinted.

PharmCast version 10 is calibrated across 74 to 1000 Da, most tightly on screening-collection chemistry and loop peptides, and least tightly on the diversity of activity-backed ChEMBL chemistry. Agreement declines at the upper end of the mass range, where the training set is thinnest. The donor, acceptor and rotatable-bond ranges are given in section 3.6.2 because a molecule can sit inside the mass range and outside the training set on any of them.

### 3.5 Performance by training-set size and composition

All released models can be compared on one curve. The plotted agreement is the median MCC, and every version is measured on the same 155,648 screening collection test cases excluded from every training set. The curve therefore compares nine models on one population.

Agreement reads 0.833 at 1.72 million molecules, 0.837 at 2.11 million, 0.840 at 2.60 million, 0.845 at 2.89 million, 0.855 at 3.28 million, 0.877 at 3.72 million, 0.879 at 4.24 million, 0.880 at 5.13 million and 0.881 at 5.89 million. Table 10 separates screening collection chemistry from real ChEMBL compounds above 600 Da. Screening collection ranking accuracy rises from 0.908 and then levels off in the high 0.93s. Large-ChEMBL ranking accuracy is 0.701 at PharmCast version 2, 0.710 at version 3 and 0.711 at version 4. Over that range the training set grew by 51%, but the added molecules came from a library filtered at 600 Da, so the large-compound result did not move.

Adding chemistry absent from the collection improved the out-of-domain result. Large-compound ranking accuracy reads 0.711 at PharmCast version 4, 0.746 at PharmCast version 5, 0.811 at PharmCast version 6, 0.832 at PharmCast version 7, 0.822 at PharmCast version 8, 0.817 at PharmCast version 9 and 0.826 at PharmCast version 10.

### 3.6 Version 10 performance

Table 7 gives the version 10 training set by source.

**Table 7.** The PharmCast version 10 training set by source. A molecular-weight-stratified one percent was held back for early stopping.

| Source | Molecules | Share | Mass range |
| --- | --- | --- | --- |
| Screening collection | 4,563,262 | 77.51% | 74 to 600 Da |
| ChEMBL, activity backed | 1,202,183 | 20.42% | 142 to 1000 Da |
| Protein loop peptides, from PDB extractions | 121,784 | 2.07% | 116 to 988 Da |
| Total | 5,887,229 | 100% | 74 to 1000 Da |

PharmCast version 10 ran for 100 epochs and restored the weights of epoch 68. Table 8 reports it on the three held-out populations. Table 10 and Figures 9 and 10 report version-to-version comparisons.

**Table 8.** PharmCast version 10 on three held-out populations. Screening collection is the Enamine test cases excluded from every training set, ChEMBL is activity-backed chemistry not present in the version 10 training set, and loop peptides are the reserved holdout. None was used for training or early stopping.

| Population | Molecules tested | Median MW | Ranking accuracy | Median error | Pearson r | Pairs within 0.05 |
| --- | --- | --- | --- | --- | --- | --- |
| Screening collection | 155,648 | 211.3 | 0.936 | 0.008 | 0.980 | 89.3% |
| Loop peptides | 13,500 | 575.7 | 0.952 | 0.016 | 0.984 | 88.3% |
| ChEMBL, activity backed | 139,700 | 386.4 | 0.889 | 0.027 | 0.936 | 74.8% |

**Table 9.** Descriptor scope of the version 10 training set, 5th to 95th percentile of a 20,000-molecule sample.

| Source | MW | MW median | Donors | Acceptors | Rotatable bonds |
| --- | --- | --- | --- | --- | --- |
| Screening collection | 256 to 467 | 344 | 0 to 3 | 2 to 7 | 2 to 8 |
| ChEMBL | 217 to 378 | 323 | 0 to 3 | 2 to 7 | 1 to 8 |
| Loop peptides | 387 to 778 | 575 | 5 to 12 | 5 to 11 | 10 to 24 |

**Table 10.**
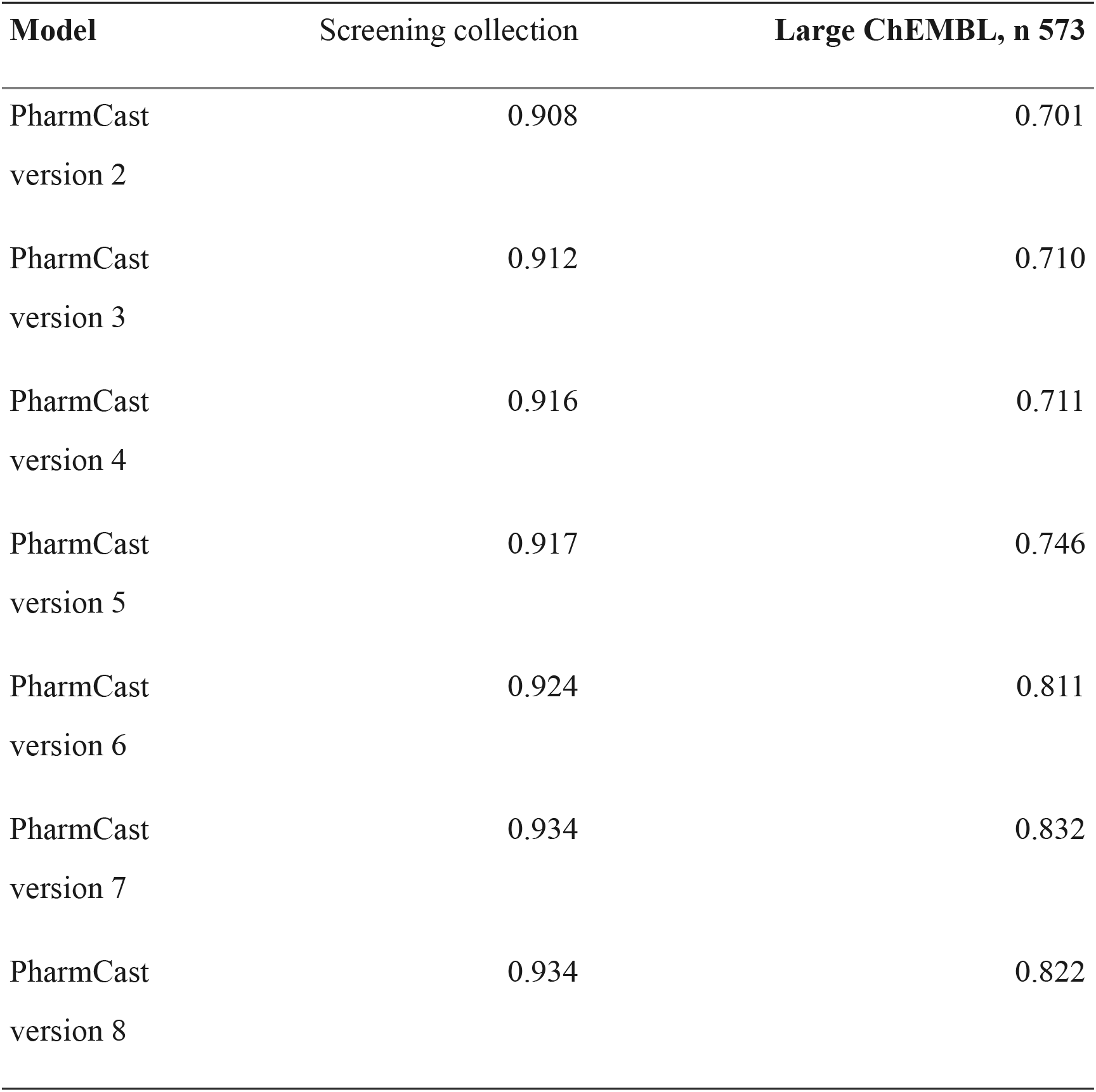
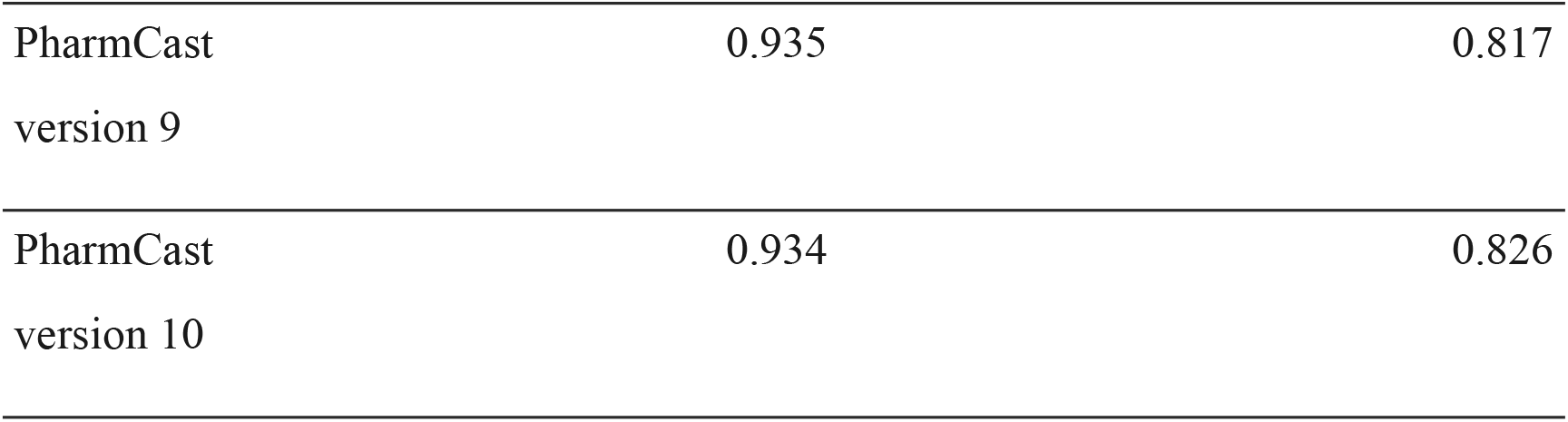
Ranking accuracy against training-set size on fixed populations held out from every model. The first population is 5,000 held-out screening collection molecules reserved from every model’s training set; large ChEMBL is 573 real compounds above 600 Da.

**Figure 9.**
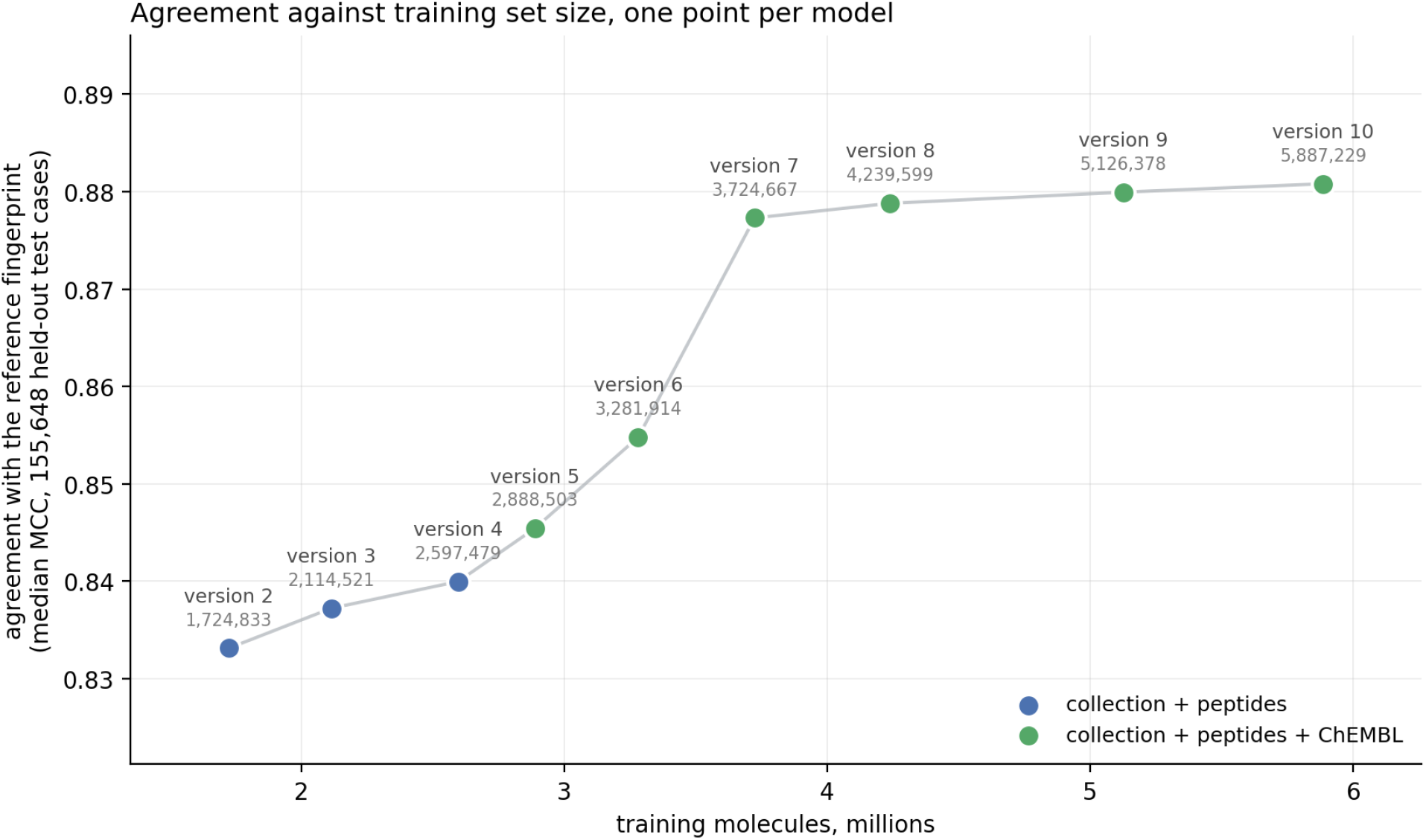
Median MCC against training-set size on all 155,648 held-out screening collection test cases. Every model is scored on the same held-out test cases, so the points differ only by the model. The test cases are held out from every version by construction. The series ends at PharmCast version 10. Color marks the training sets; each point is labeled with its training set size.

**Figure 10.**
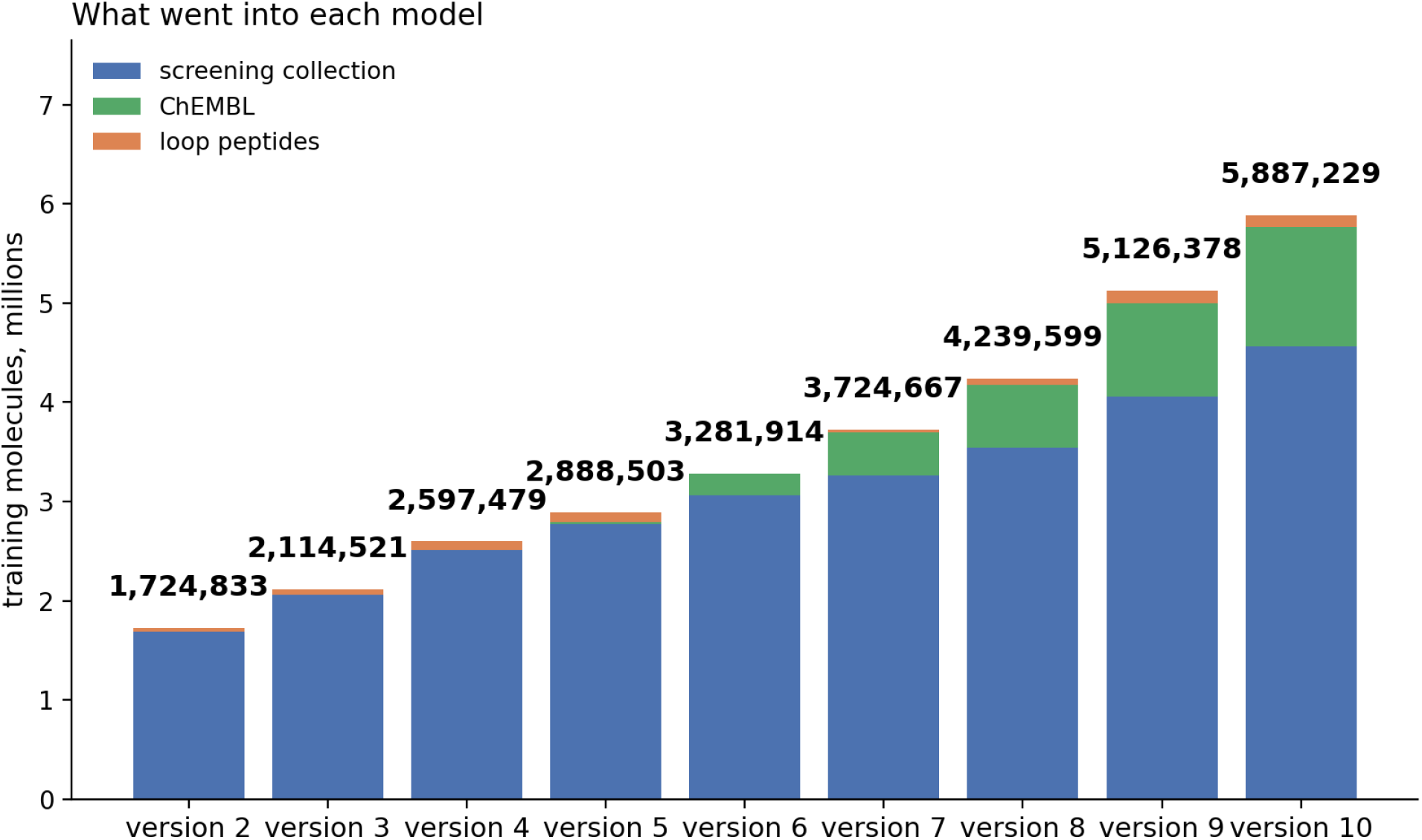
Training-set composition and total count across the three source sets for each model. PharmCast version 4 is the last model without ChEMBL; PharmCast version 5 was the first to include all three source sets; the series ends at PharmCast version 10.

The source sets continue to be fingerprinted as they grow. Subsequent models can incorporate the expanded sets.

#### 3.6.1 Training-set composition

Figure 11 and Table 1 report the composition of the version 10 training set. The screening collection contributes 4,609,488 compounds and is 99.8% complete. The loop-peptide set is complete at 136,494. ChEMBL contributes 1,214,214 compounds and is 44.4% complete. Additional eligible ChEMBL compounds may extend these ranges in later versions.

**Figure 11.**
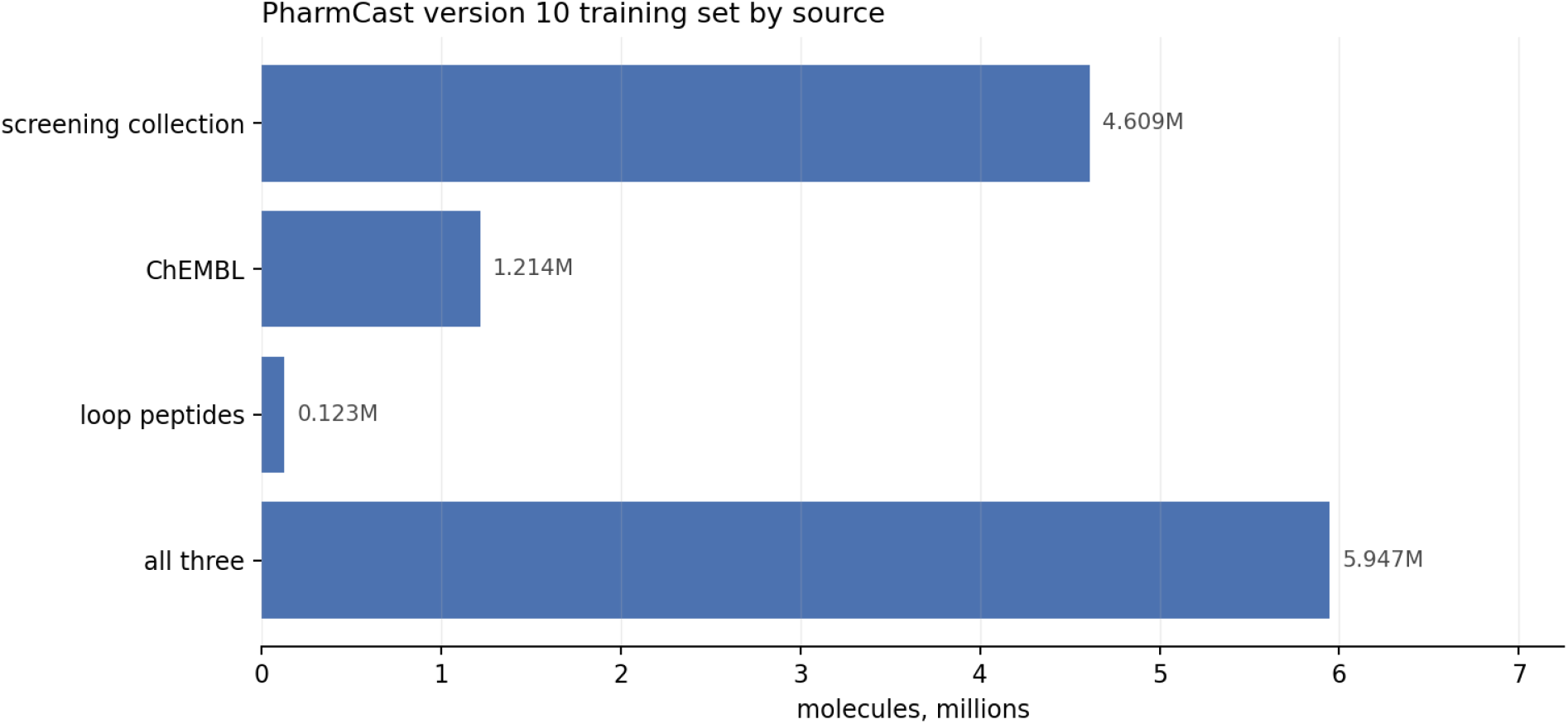
The PharmCast version 10 training set by source. The screening collection contributes 4.609 million molecules, ChEMBL 1.214 million and the loop peptides 0.123 million, giving 5.947 million in all.

#### 3.6.2 Descriptor coverage and historical performance

Section 3.4 reports the mass axis, on which this model’s error moves, but mass is not the only axis. Donors and acceptors are two of the six feature types the fingerprint is built from, and the rotatable bond count sets how much conformational space the ensemble has to cover, so a molecule can sit inside the mass range and outside the training set on either. Each range below is the 5th to the 95th percentile of a random sample of 20,000 molecules per source, with the full observed range beneath.

Full observed ranges over the same samples are as follows: the collection spans 173 to 597 Da, 0 to 5 donors, 0 to 11 acceptors and 0 to 10 rotatable bonds; ChEMBL spans 142 to 646 Da, 0 to 12 donors, 0 to 19 acceptors and 0 to 27 rotatable bonds; and the peptides span 159 to 961 Da, 0 to 19 donors, 2 to 13 acceptors and 2 to 32 rotatable bonds. The collection and peptide rows describe their full source sets. The ChEMBL row describes the reference fingerprints used for version 10; additional eligible ChEMBL compounds may extend these ranges in later versions. The rotatable-bond distribution separates the three source sets more strongly than molecular weight: median values are 5 for the screening collection, 4 for the ChEMBL set in the version 10 training set, and 16 for the peptides.

Table 10 tracks ranking accuracy against training-set size for every released model from v2 onward. Each model is scored on the same 5,000 Enamine screening collection test cases and the same 573 real ChEMBL compounds above 600 Da. Both populations were excluded from every model’s training set.

### 3.7 Application: screening collection search against a pharmacophore reference

A fast pharmacophore comparison can be used as a search objective. Orforglipron, an 883 Da non-peptidic reference, was screened against 4,653,831 collection compounds, with a fingerprint generated from SMILES for each, in 29.3 minutes. Screening the same collection when the fingerprints already exist is a bitwise comparison at 0.012 ms per pair. The leading compounds were then recomputed with the reference calculation. The highest-scoring purchasable compound has an ensemble Tanimoto of 0.786 at 591 Da and a two-dimensional Morgan similarity of 0.113 to the reference, supporting scaffold hopping instead of analog retrieval.

The reference value of 0.786 establishes a baseline that subsequent designed molecules must exceed. This application also lies in a sparsely populated region of the training set, motivating the training set expansion reported here.

## 4. Discussion

PharmCast predicts an ensemble pharmacophore fingerprint from two-dimensional structure with median error of 0.008 on screening collection chemistry and 0.016 on conformationally rich peptides, against 0.006 for the reference calculation’s own reproducibility. Because the fingerprint is an OR over conformations, it records what a molecule can present without committing to a particular pose. This is primarily a property of molecular constitution, making the task more tractable than conformer prediction.

Error is governed by training-set coverage, and we expect this relationship to generalize beyond PharmCast. The historical comparisons are confined to Figures 9 and 10, Table 10 and the paragraphs that read them directly. The difference was training-set composition. Additional screening collection data filtered at 600 Da did not improve the large-compound population, whereas adding ChEMBL chemistry did.

Coverage is not a proxy for size. Section 3.4 measures agreement against molecular weight and finds it flat across screening collection chemistry, and lowest on ChEMBL compounds below 400 Da, which are small. A single accuracy figure over a pooled evaluation set would have concealed all of this.

These results support reporting performance stratified by the property associated with failure and distributing the applicability domain with each model. The model card records the mass range, training-set composition, and measured degradation so that users can evaluate the boundary without consulting this paper.

The peptide set may also prove useful for more than training. Its molecules are real loops taken from deposited structures, conformationally rich in a way screening collection chemistry is not, and each one is a presentation some protein was observed to hold, which makes them a natural reference set for peptide mimicry, where a small molecule is scored against the pharmacophoric presentation of a loop instead of against another small molecule.

When predicted similarity is the objective of a search, the compounds that rise to the top include those the model overestimates, so the scores of the winners are optimistic. The remedy is to let the surrogate rank and the reference calculation decide. In a CHIP de novo design campaign against orforglipron, PharmCast ranked every candidate and the survivors were then recomputed with the reference calculation, so each similarity reported for that campaign is a measurement rather than a prediction: the delivered design matches the reference at 0.810 pharmacophore similarity while sharing 0.145 two-dimensional similarity with it, and in the refinement stage the best 250 candidates by PharmCast were rescored, the best of them measuring 0.797 against the parent’s 0.754. We therefore recommend ranking with the surrogate, deciding with the reference calculation, and carrying a broad set of survivors into the rescoring step, so that no choice is made on a predicted score.

## 5. Limitations

The model predicts the ORed ensemble fingerprint. Where an application requires the matching geometry of a particular conformer, that must come from the reference calculation, which retains per conformer records; the surrogate stands in for the ensemble, not for the conformers behind it.

Performance remains lowest outside screening-collection chemistry, particularly for the diversity of activity-backed ChEMBL compounds. Section 3.4 attributes this degradation to limited training-set coverage. Above 700 Da, the training set is almost entirely peptidic, and agreement decreases.

Predicted similarities have a small positive bias caused by systematic overprediction of set bits, with a median signed error of +0.002 on screening collection chemistry. This bias does not affect ranking but should be considered when applying absolute thresholds.

The screening collection has a median nearest-neighbor similarity of 0.714, so held-out molecules drawn from it are not fully independent of the version 10 training set.^24^ For the version 10 screening collection test set, half of the held-out split is within 0.691 of a training molecule, and agreement increases from 0.857 for molecules in the 0.35 to 0.50 band to 0.922 in the 0.70 to 1.00 band. The in-domain values therefore describe performance on chemistry of the same kind, not on unseen chemistry. The peptide set and the collection differ in both size and character, so the peptide result and the collection result are not evidence about each other.

All results reported here are computed on real screening collection compounds, on peptides extracted from experimentally determined structures, and on real ChEMBL entries. No molecules were generated for evaluation.

## 6. Author contributions

S.M.M. conceived the study, constructed the source sets and surrogate, performed the evaluation, and wrote the manuscript. M.J.M. co-developed the pharmacophore fingerprinting method on which this work builds and contributed to the analysis and manuscript.

## 8. Data and code availability

The PharmCast package, model utilities, and documentation are available at https://github.com/smuskal/pharmcast under Apache-2.0, which covers both code and model weights. Trained weights are distributed from https://pharmcast.ai. CHIP is the de novo design protocol introduced in reference 25 and since refined; the campaign reported in the Discussion is described at https://www.eidogen.com/chip2026_metoo.php.

The repository does not include the reference pharmacophore fingerprint generator or its toolchain and does not redistribute the training sets. Screening collection and building-block data are supplied under Enamine’s terms. ChEMBL-derived data are licensed under CC BY-SA 3.0.

Each model checkpoint records its training-set composition, mass range, and applicability domain. These values can be displayed with the PharmCast model-card command applied to M.pt.

